# The role of recombination dynamics in shaping signatures of direct and indirect selection across the *Ficedula* flycatcher genome

**DOI:** 10.1101/2022.08.11.503468

**Authors:** Madeline A. Chase, Carina F. Mugal

## Abstract

Recombination is a central evolutionary process that reshuffles combinations of alleles along chromosomes, and consequently is expected to influence the efficacy of direct selection via Hill-Robertson interference. Additionally, the indirect effects of selection on neutral genetic diversity are expected to show a negative relationship with recombination rate, as background selection and genetic hitchhiking are stronger when recombination rate is low. However, owing to the limited availability of recombination rate estimates across divergent species, less is known about the impact of evolutionary changes in recombination rate on genomic signatures of selection. To address this question, we estimate recombination rate in two *Ficedula* flycatcher species, the taiga flycatcher (*F. albicilla*) and collared flycatcher (*F. albicollis*). We show that recombination rate is strongly correlated with signatures of indirect selection, and that evolutionary changes in recombination rate between species have observable impacts on this relationship. Conversely, signatures of direct selection on coding sequences show little to no relationship with recombination rate, even when restricted to genes where recombination rate is conserved between species. Thus, using measures of indirect and direct selection that bridge micro- and macro-evolutionary timescales, we demonstrate that the role of recombination rate and its dynamics varies for different signatures of selection.

## INTRODUCTION

Meiotic recombination is a central evolutionary process, which can influence genome evolution via a range of different mechanisms. On the one hand, recombination may be an evolutionary advantage contributing to the origin of sexual reproduction (1-3). By reshuffling variation, recombination can create novel combinations of alleles, which can aid in adaptation. However, recombination may not be universally advantageous. The same reshuffling that may create adaptive combinations of alleles can also break apart existing adaptive associations. In some systems, there is evidence that the evolution of suppressed recombination can promote local adaptation and speciation (4-8), for instance when inversions capture multiple loci with beneficial variation. In addition to breaking up linkage among sites, recombination is associated with the process of GC-biased gene conversion (gBGC), which leads to the preferential fixation of G:C over A:T alleles and can interfere with fitness (9, 10). Thus, recombination can have a multi-faceted impact on fitness.

Variation in recombination rate across the genome can also play a major role in shaping genomic signatures of natural selection. Recombination rate can impact the efficacy of selection by Hill-Robertson interference (HRI), where linkage between multiple non-neutral mutations leads to selective interference (11). Consequently, HRI predicts that the efficacy of natural selection should increase with increasing recombination rate, reflected in signatures of direct selection by increased fixation of deleterious mutations where recombination rate is low and increased fixation of beneficial mutations where recombination rate is high. Typical measures of direct selection used to assess the presence of HRI are the nonsynonymous over synonymous ratio of diversity π_N_/π_S_, and the nonsynonymous over synonymous ratio of divergence *d*_*N*_*/d*_*S*_, which are predicted to show a negative relationship with recombination rate, and the adaptive substitution rate ω_a_, which is predicted to show a positive relationship with recombination rate (Fig. 1C). Besides influencing the efficacy of selection, physical linkage among sites also manifests in a reduction of genetic diversity at neutral sites that are linked to targets of selection. Physical linkage creates local reductions in neutral diversity in response to selective sweeps through a process called genetic hitchhiking (HH; 12). The size of a selective sweep depends not only on the strength of selection, but also on the recombination rate, with population genetic theory showing that the reduction in diversity rapidly decreases with increasing recombination (13,14). This means selective sweep signatures are predicted to correlate negatively with recombination rate (Fig.1C). In addition to HH, background selection (BGS) reduces diversity at sites linked to deleterious mutations with a greater impact where recombination rate is lower (15). We will jointly refer to HH and BGS as indirect selection, due to the key feature that they refer to the indirect effects of selection on linked, neutral sites. The diversity reducing effects of indirect selection are thought to contribute to driving elevated levels of genetic differentiation (*F*_*ST*_) between species (16,17), resulting in a negative relationship between *F*_*ST*_ and recombination rate (Fig. 1C).

**Figure 1:**
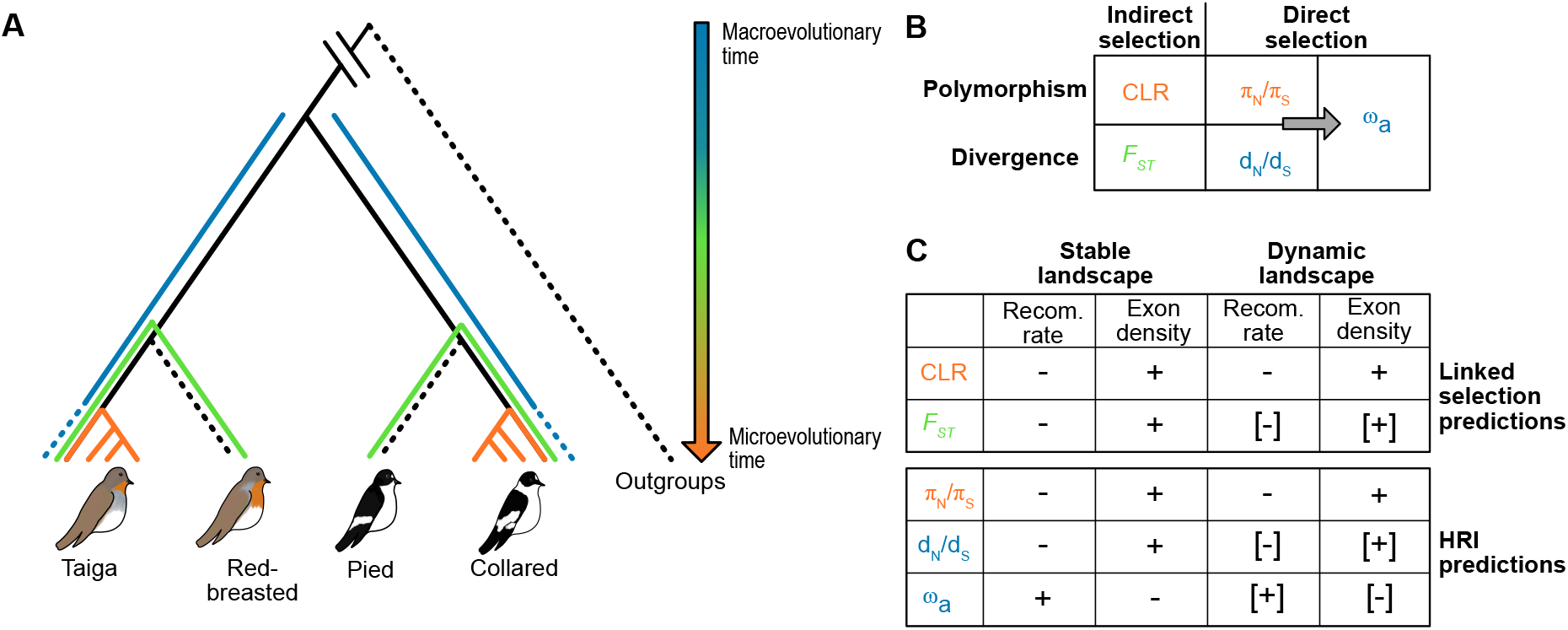
Outline of the study design and working hypothesis. Shown in panel (A) is the topology of the species included in the present study. In colour we indicate the timescale, from a macroevolutionary (blue) to a microevolutionary (orange) timescale, of different measures of natural selection estimated in the present study and listed in panel (B). The two polymorphism-based measures, the composite likelihood ratio (CLR) test for selective sweeps and π_N_/π_S_ are shown in orange and are estimated for taiga flycatcher and collared flycatcher. *F*_*ST*_ is shown in green and is estimated between taiga and red-breasted flycatcher and between collared and pied flycatcher. Lineage-specific *d*_*N*_*/d*_*S*_ is shown in blue and is estimated for taiga flycatcher and collared flycatcher with help of an outgroup species. The dashed blue line indicates that estimates of *d*_*N*_*/d*_*S*_ are corrected for within-species polymorphisms. Species that were only used as outgroups or as reference species for estimating pair-wise *F*_*ST*_ are represented by dashed black lines. Panel (C) summarizes the predicted relationships between different measures of selection and recombination rate and density of selected sites in the presence of linked selection and HRI. Predictions are presented for a stable recombination landscape versus a dynamic recombination landscape, where correlations are shown with brackets if we expect them to be weakened by a dynamic recombination rate.

Many population and speciation genomic studies have investigated the relationship between recombination rate and genomic signatures of indirect selection. Across a wide range of taxa, variation in genetic diversity is strongly correlated with recombination rate (18-20). Differentiation islands, genomic regions showing significantly elevated *F*_ST_ relative to the genomic background, also frequently correspond to regions of reduced recombination (21-26). Additionally, in species with ongoing gene flow, barrier loci and reductions in introgression have been observed in regions of low recombination (27-30). Altogether, these observations point to a clear and pervasive role of recombination rate shaping signatures of indirect selection and correspond well with the expectations from population genetic theory.

On the other hand, evidence for recombination rate shaping signatures of direct selection through HRI appears less consistent across different taxa. HRI has been suggested to play a role in shaping patterns of molecular evolution in many systems including, among others, the invertebrates *Drosophila* (*D*.) *melanogaster* (31-33), *D. pseudoobscura* (34), *Heliconius melpomene* (35), the vertebrate great tit (*Parus major*) (36), and the plant sorrel (*Rumex hastatulus*) (37). However, in other systems, including collared flycatcher (*Ficedula* (*F*.) *albicollis*), there has been no evidence of HRI shaping the efficacy of selection across the genome (38-41). One potential explanation for mixed evidence for a relationship between recombination rate and signatures of direct selection is that the intensity of HRI relies on multiple interacting variables, which may differ greatly among divergent taxa. For example, the density of functional sites and mutation rate contribute to the strength of HRI because interference requires multiple selected mutations to be linked and segregating at the same time. The addition of neutral sites in between selected sites (i.e. introns in between exons) can lessen the impact of interference (42, 43). Thus, the impact of HRI on measures of direct selection may be more apparent in more compact genomes, such as invertebrate genomes, where gene density is higher compared to vertebrates and plants. Additionally, if there is rapid evolution of the recombination landscape, then a relationship between recombination rate and genomic signatures of direct selection may not have the chance to build up (34).

The dynamics of recombination rate evolution are highly variable across different taxa. For instance, in many mammals, a rapid turnover of recombination hotspots is observed. The fast evolution of hotspots is mediated by the zinc-finger protein PRDM9 (44-46), which encodes the location of hotspots and is rapidly evolving in response to the erosion of hotspot motifs (47-49). In organisms that lack PRDM9, such as birds, recombination hotspots tend to be more stable over evolutionary time and the recombination landscape evolves at a slower pace (50, 51). In addition to recombination hotspot evolution, also broad-scale variation in recombination rate changes over time, for example, as a result of chromosomal rearrangements (52-55). However, despite a broad acknowledgement that recombination rate is dynamic, only few studies have directly investigated the impact of recombination rate evolution on genomic signatures of natural selection.

Here we investigate the role of recombination rate dynamics in shaping patterns of indirect and direct selection in two species of *Ficedula* flycatchers (Fig. 1A). For this purpose, we estimate lineage-specific recombination rates in both species, as well as four measures of selection, with two measures each of indirect and direct selection, a polymorphism-based and a divergence-based measure, respectively (Fig. 1); these are the composite likelihood ratio (CLR) test for selective sweeps, genetic differentiation *F*_*ST*_, π_N_/π_S_, and *d*_*N*_*/d*_*S*_ (Fig. 1B). Additionally, we estimate the adaptive substitution rate (ω_a_), which relies on a combination of polymorphism and divergence data. We then investigate the relationship between evolutionary changes in recombination rate and these measures of selection. We also study the association between selective sweeps and the adaptive substitution rate, i.e. two distinct genomic signatures of positive selection, which respectively assess indirect and direct selection. In brief, we address the following questions: 1) How does recombination rate impact genomic signatures of indirect and direct selection in *Ficedula* flycatchers? 2) How do the dynamics of recombination rate evolution impact both signatures of selection? and 3) How do signatures of indirect and direct selection compare to one another?

## RESULTS

### Study system and whole genome re-sequencing data

To understand the relationship between the evolutionary dynamics of recombination rate and genomic signatures of indirect and direct selection, we collated whole genome re-sequencing data for four species of *Ficedula* flycatchers: 65 taiga flycatchers (*F. albicilla*; 56), 15 red-breasted flycatchers (*F. parva;*(56)), 95 collared flycatchers (*F. albicollis*; 57), and 11 pied flycatchers (*F. hypoleuca*; 21). Additionally, we included one individual of snowy-browed flycatcher (*F. hyperythra*; 21) as an outgroup for variant polarization. We performed variant calling on all five flycatcher species and identified in total 51,424,863 single nucleotide variants (SNVs) within a set of 566,724,393 callable sites. We then focus our study on the more distant comparison of taiga flycatcher and collared flycatcher, while red-breasted flycatcher and pied flycatcher are only included for SNV polarization and to estimate pairwise *F*_*ST*_ (Figure 1).

### The evolutionary dynamics of recombination rate among taiga and collared flycatchers

To assess the evolutionary dynamics of the recombination rate in *Ficedula* flycatchers, we first estimated recombination rate in taiga flycatcher and collared flycatcher based on patterns of linkage disequilibrium (LD). LD-based estimates of recombination rate have been shown to be influenced by recent demography, where in particular fine-scale variation and recombination hotspot detection appears to be affected (58) However, simulation results suggest that broad-scale estimates of recombination rate are more robust to the demographic history (59). We therefore focus on broad-scale recombination estimates at a resolution of 200-kb windows. LD-based estimates for both taiga flycatcher and collared flycatcher showed significant correlations with earlier pedigree-based estimates for collared flycatcher (60), with Pearson’s correlation coefficients *R* = 0.46 and *R* = 0.48, *p*-value < 10^−16^,respectively. We then used the collared flycatcher linkage map to convert LD-based recombination rate from population-scaled recombination (*ρ* = 4*N*_*e*_r) into cM/Mb (see Materials and Methods).

We observed a significant correlation for recombination rate between the two species estimated in 200-kb windows (Fig. S1; Pearson’s *R* = 0.45, *p*-value < 10^−16^). The correlation here is weaker than correlations observed previously between the more closely related collared and pied flycatchers (51; Pearson’s *R* = 0.79). The observed differences in recombination rate between taiga flycatcher and collared flycatcher allow us to investigate how changes in recombination rate impact different forms of selection.

### Recombination rate shapes genomic signatures of indirect selection

We examined the relationship between recombination rate and measures of indirect selection, for which we computed two statistics, the CLR test for selective sweeps and pairwise *F*_*ST*_ between the species pairs taiga and red-breasted flycatcher and collared and pied flycatcher (Fig. 1A). Logistic regression analysis between lineage-specific recombination rate and CLR estimates in both taiga flycatcher and collared flycatcher support the predicted relationship between recombination and selective sweep signatures (Table 1; Fig. S2), with selective sweeps tending to occur in windows with lower recombination rate. To investigate how evolutionary changes in recombination rate impact genomic signatures of indirect selection, we performed logistic regression analysis with recombination rate estimated in the other species. This revealed that the relationship was weaker when we compared selective sweep signatures in one species with recombination rate in the other species (Table 1; Fig. S2).

**Table 1:** The relationship between recombination rate and selective sweep presence. Shown are the results of logistic regression analyses between presence/absence of a selective sweep and LD-based recombination rate in 200-kb windows; expected log odds estimate, standard error (SE), Z-standardized log odds estimate (Z-value), the McFadden *R*^2^, and *p-*value. Regression analyses were performed using sweep presence/absence and recombination rate estimated in the same species, as well as recombination rate in the other species.

| Sweeps/recombination | Estimate | SE | Z-value | $R^2$ | $p$-value |
| --- | --- | --- | --- | --- | --- |
| taiga/taiga | -2.6 | 0.18 | -14 | 0.38 | $< 2.2 \cdot 10^{-16}$ |
| collared/collared | -6.3 | 1.0 | -6.1 | 0.62 | $8.8 \cdot 10^{-10}$ |
| taiga/collared | -0.39 | 0.063 | -6.1 | 0.027 | $1.0 \cdot 10^{-9}$ |
| collared/taiga | -0.34 | 0.11 | -3.0 | 0.028 | $2.3 \cdot 10^{-3}$ |

Next, we compared how lineage-specific estimates of recombination rate corresponded with the differentiation landscape for two species comparisons. We observed a negative relationship between recombination rate and *F*_*ST*_ for both species pairs, red-breasted and taiga flycatchers (Fig. S3; Spearman’s *R* = -0.71; *p*-value < 2.2·10^−16^) and collared and pied flycatchers (Fig. S3; Spearman’s *R* = -0.59; *p*-value < 2.2·10^−16^), which is in good agreement with the linked-selection prediction. Comparing the distribution of recombination rate in different categories of *F*_*ST*_ peak revealed that both taiga flycatcher and collared flycatcher show a significant reduction in recombination rate for shared *F*_*ST*_ peaks (Fig. 2) compared to windows without *F*_*ST*_ peaks. Collared flycatcher showed a significant reduction in recombination rate in collared/pied-specific (CP unique) *F*_*ST*_ peaks, but not for red-breasted/taiga-specific (TR unique) peaks (Fig. 2B). Although taiga flycatcher showed a significant reduction in recombination rate for both TR unique peaks as well as CP unique peaks compared to the background without peaks, the reduction in recombination rate was greater for shared and TR unique peaks compared to CP unique peaks (Fig. 2A). This supports previous evidence that suggested that changes in recombination rate underlie lineage-specific *F*_*ST*_ peaks in *Ficedula* flycatchers (56). Moreover, these results demonstrate that not only are measures of indirect selection strongly impacted by recombination rate, but that also the evolutionary dynamics of recombination rate play a crucial role.

**Figure 2:**
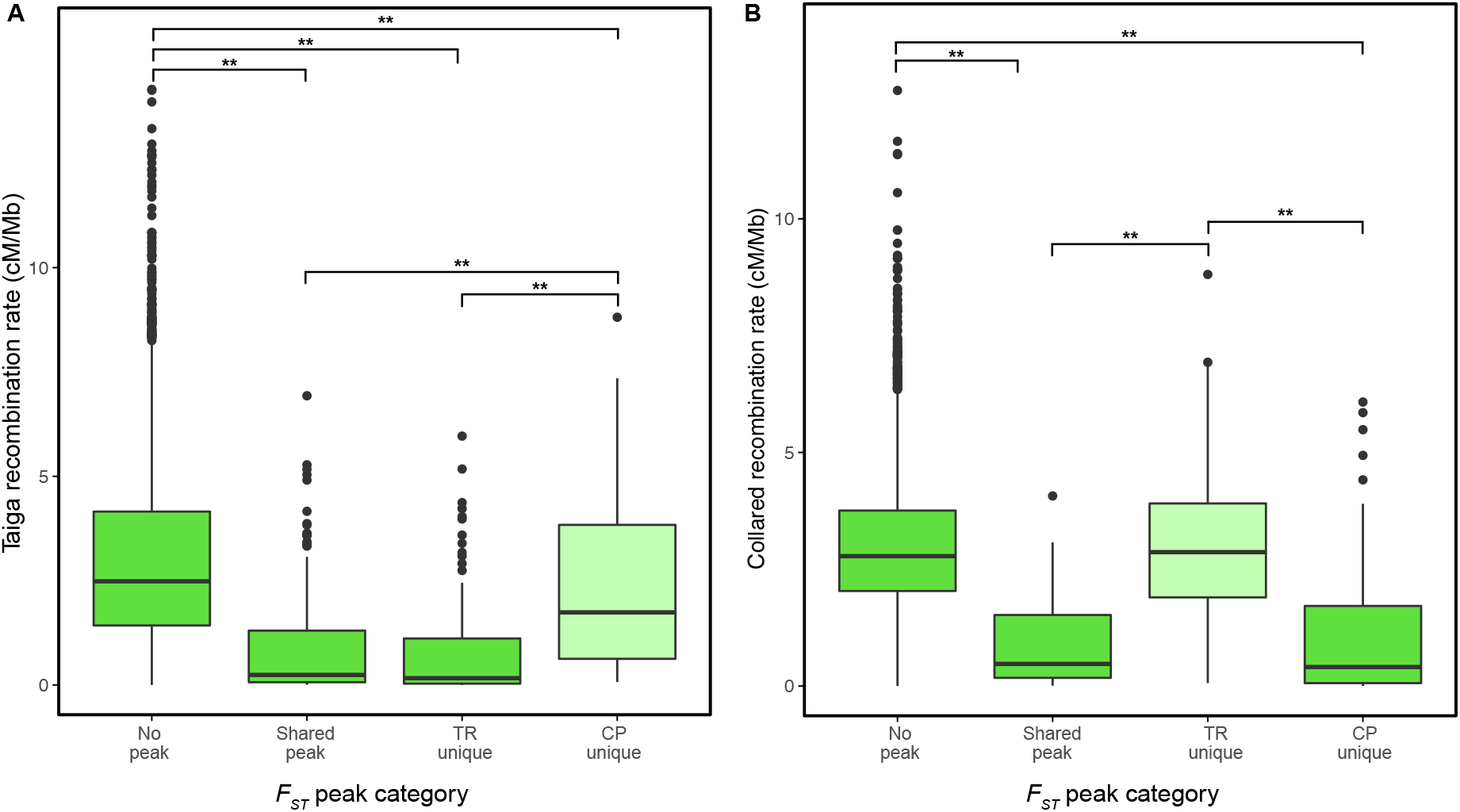
The impact of recombination rate on indirect estimates of selection. Shown is the relationship between lineage-specific recombination rate and window-based estimates of *F*_*ST*_ for both (A) taiga flycatcher and (B) collared flycatcher. The 200-kb windows are divided by windows with no *F*_*ST*_ peaks, *F*_*ST*_ peaks shared between both species pairs, and two categories of lineage-specific *F*_*ST*_ peaks, i.e. taiga/red-breasted-specific *F*_*ST*_ peaks (TR unique) or a collared/pied-specific *F*_*ST*_ peaks (CP unique). Asterisks show significance based on 1000 permutations, * *p*-value <0.05, ** *p*-value <0.01. See Table S1 for the number of windows in each category.

### Genomic signatures of direct selection are not consistent with HRI

We examined the relationship between recombination rate and lineage-specific signatures of direct selection, π_N_/π_S_, *d*_N_/*d*_S_, and ω_a_ across genes overlapping with three approximately equal-sized recombination rate bins, referred to as low (L), intermediate (M), and high (H). Since we might expect that any relationship between recombination rate and measures of direct selection would be weakened by evolutionary changes in recombination rate, we also estimated π_N_/π_S_, *d*_N_/*d*_S_, and ω_a_ only for genes where recombination rate is conserved between taiga flycatcher and collared flycatcher (See Materials and Methods). In addition, we examined the distribution of fitness effects (DFE) and in particular the mean strength of selection against deleterious mutations *N*_e_*s*, since recent work has shown that changes in population size can create spurious correlations between ω_a_ and any variable that itself is correlated to the parameter *N*_e_*s* (61). We note, however, that it has been shown that estimates of *N*_e_*s* can vary widely, sometimes taking very extreme values (62). Finally, to account for any potential bias due to gBGC, estimates of π_N_/π_S_, *N*_e_*s, d*_N_/*d*_S_, and ω_a_ are based on GC-conservative changes only. Results based on all changes are shown in the Supplement.

Generally, we did not observe the relationships between recombination rate and signatures of direct selection predicted by HRI in either species (Fig. 3), regardless of whether we restricted the analysis to genes with conserved recombination rate or not. None of the statistics showed a significant difference in any of the recombination rate bins for taiga flycatcher. Collared flycatcher showed a slight negative trend in π_N_/π_S_ and a slight positive trend in *N*_e_*s* as recombination rate increases, indicative of an increase in the efficacy of purifying selection with increasing recombination rate, although the differences were not statistically significant. However, *d*_N_/*d*_S_ and ω_a_ were both highest for the intermediate recombination rate bin, a pattern also observed previously (63) and not consistent with HRI.

**Figure 3:**
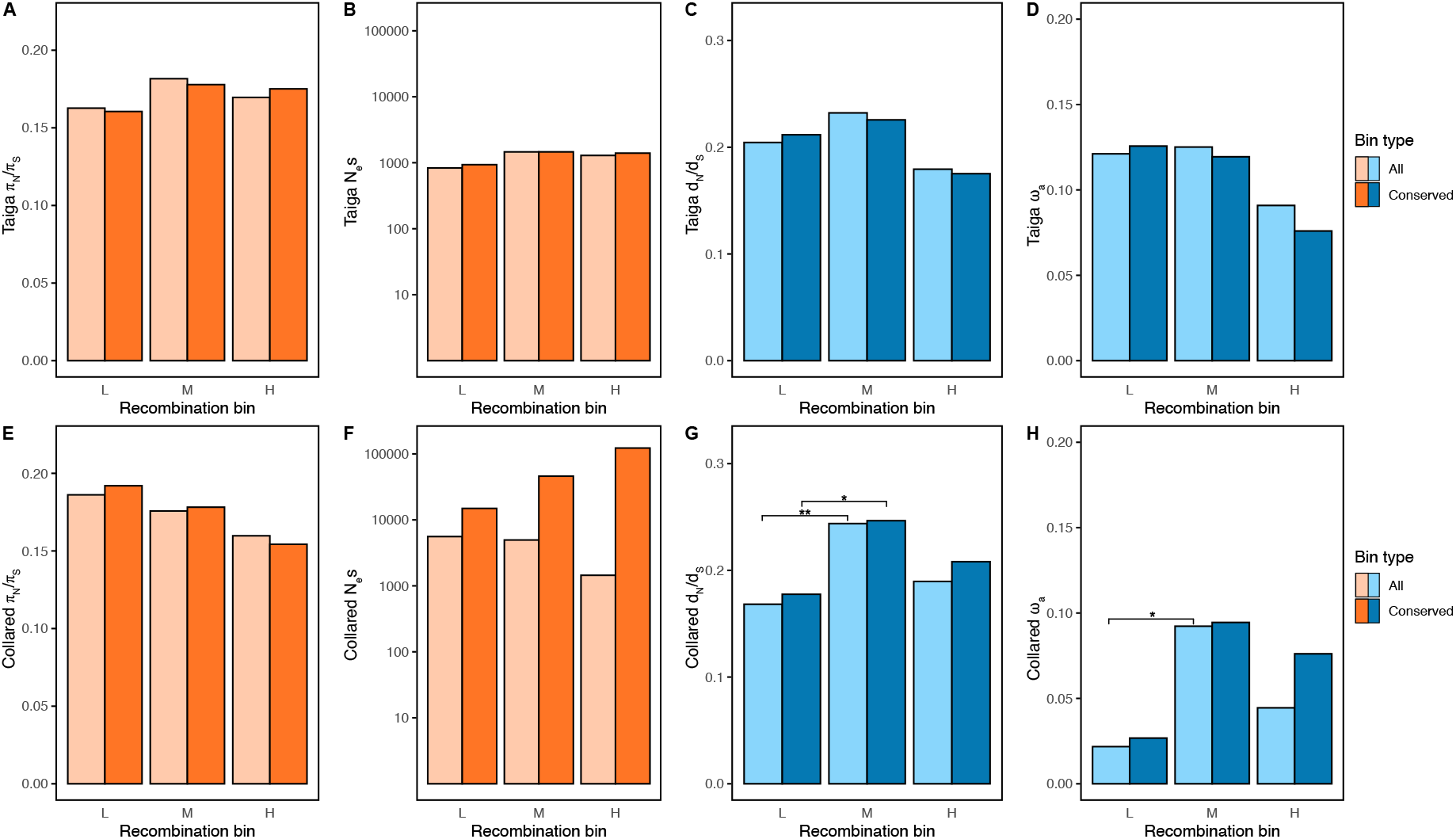
The impact of recombination rate on measures of direct selection. Shown are estimates of π_N_/π_S_, *N*_e_*s, d*_N_/*d*_S_ and ω_a_ for (A-D) taiga flycatcher and for (E-H) collared flycatcher estimated for genes overlapping with 3 bins of recombination rate (low (L), intermediate (M) and high (H)). Genes are separated based on conservation of recombination rate in taiga flycatcher and collared flycatcher. Colours for bin type correspond to whether the estimate shown is calculated for all genes (lighter) or for genes overlapping with windows with conserved recombination rate (darker). Tests for significance were performed between recombination rate within bin types (all or conserved). Asterisks denote statistical significance, * *p*-value <0.05; ** *p*-value <0.01. See Fig. S4 for plots based on all changes. See Table S2 for the number of genes in each bin.

To examine whether variation in the density of selected sites might obscure patterns of HRI, we divided the recombination rate bins into low and high functional density (i.e. density of exons and conserved non-coding elements; CNEs). We observed no consistent pattern between species that matches the predictions based on HRI for recombination rate and signatures of direct selection (Fig. 4). Taken together, these results suggest a negligible role of recombination in shaping genomic signatures of direct selection in *Ficedula* flycatchers.

**Figure 4:**
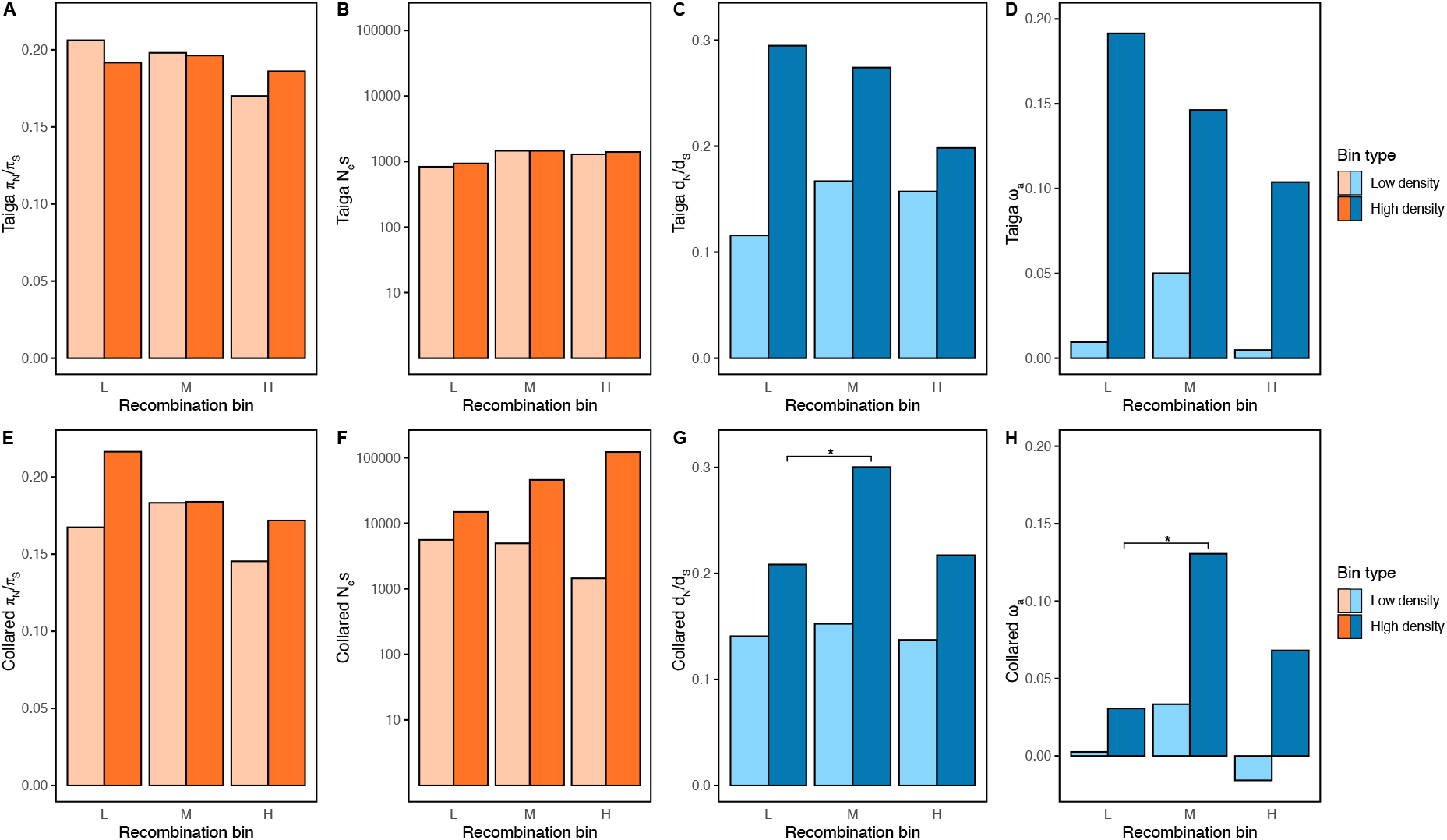
The impact of gene density on measures of direct selection. Shown are estimates of π_N_/π_S_, *N*_e_*s, d*_N_/*d*_S_ and ω_a_ for (A-D) taiga flycatcher and for (E-H) collared flycatcher estimated for genes overlapping with 3 bins of recombination rate (low (L), intermediate (M) and high (H)). Genes are separated based on whether they are found in windows with low or high functional density. Tests for significance were performed between recombination bins within both bin density types. Asterisks denote statistical significance based on 1000 permutations, * *p*-value < 0.05, ** *p*-value <0.01. See Fig. S5 for plots based on all changes. See Table S3 for the number of genes in each bin.

### Weak association between signatures of direct and indirect selection

Finally, we compared the genomic signatures of direct and indirect selection to address the hypothesis that recurrent selective sweeps may contribute to *F*_*ST*_ peaks, particularly shared *F*_*ST*_ peaks, reflected by an increased rate of adaptive substitutions in *F*_*ST*_ peaks compared to the genomic background. For this purpose, we compared estimates of π_N_/π_S_, *N*_e_*s, d*_N_/*d*_S_, and ω_a_ for genes overlapping with shared *F*_*ST*_ peaks, lineage-specific *F*_*ST*_ peaks, and *F*_*ST*_ peaks with or without a selective sweep signature against genes not overlapping with *F*_*ST*_ peaks as a background reference. Lineage-specific *F*_*ST*_ peaks and *F*_*ST*_ peaks without a selective sweep signature showed no significant differences in signatures of direct selection compared to the genomic background. Both taiga flycatcher and collared flycatcher showed significantly higher π_N_/π_S_ in shared *F*_*ST*_ peaks relative to the genomic background (Table 2). Although *d*_N_/*d*_S_ was elevated in both species and ω_a_ elevated for taiga flycatcher, these differences were not statistically significant. Similarly, π_N_/π_S_ was significantly higher for both species within *F*_*ST*_ peaks overlapping with selective sweeps, while differences in *d*_N_/*d*_S_ and ω_a_ were not statistically significant (Table 2).

**Table 2:** Comparison of measures of indirect and direct selection. Shown are estimates of the efficacy of direct selection for genes grouped by *F*_*ST*_ peak category. Categories include genes outside *F*_*ST*_ peaks, genes in *F*_*ST*_ peaks shared between species comparisons, genes within peaks specific to the focal species, genes within peaks that show a signature of a selective sweep, and genes within peaks that do not show a signature of a selective sweep. *p*-values comparing whether statistics within *F*_*ST*_ peaks are significantly different from outside *F*_*ST*_ peaks, are presented in parentheses. Significant differences are indicated in bold, *p*-value < 0.05. See Table S4 for estimates based on all changes.

|  | Taiga flycatcher |  |  |  |  | Collared flycatcher |  |  |  |  |
| --- | --- | --- | --- | --- | --- | --- | --- | --- | --- | --- |
| | $\pi_N/\pi_S$ | $N_e s$ | $d_N/d_S$ | $\omega_a$ | genes | $\pi_N/\pi_S$ | $N_e s$ | $d_N/d_S$ | $\omega_a$ | genes |
| **Outside peaks** | 0.17 | 1276 | 0.21 | 0.11 | 5937 | 0.17 | 20817 | 0.19 | 0.051 | 5916 |
| **Shared peaks** | **0.36**<br>**(0.001)** | 113<br>(0.16) | 0.34<br>(0.059) | 0.21<br>(0.13) | 220 | **0.30**<br>**(0.009)** | 4789<br>(0.71) | 0.25<br>(0.43) | 0.017<br>(0.68) | 220 |
| **Lineage specific peaks** | 0.12<br>(0.15) | 181<br>(0.21) | 0.19<br>(0.79) | 0.16<br>(0.49) | 197 | 0.14<br>(0.63) | 154<br>(0.49) | 0.17<br>(0.73) | 0.099<br>(0.59) | 98 |
| **Peaks with sweep overlap** | **0.28**<br>**(0.004)** | 99<br>(0.14) | 0.32<br>(0.051) | 0.22<br>(0.055) | 321 | **0.33**<br>**(0.002)** | 228<br>(0.49) | 0.18<br>(0.83) | -0.065<br>(0.13) | 211 |
| **Peaks with no sweep overlap** | 0.20<br>(0.56) | 327<br>(0.34) | 0.13<br>(0.39) | 0.049<br>(0.48) | 96 | 0.17<br>(0.94) | 4732<br>(0.73) | 0.31<br>(0.17) | 0.20<br>(0.14) | 107 |

## DISCUSSION

Here we investigate the relationships between recombination rate and genomic signatures of indirect and direct selection, and how the evolutionary dynamics of recombination rate between divergent species impacts these different selection signatures. This reveals that recombination rate has a markedly more pronounced relationship with signatures of indirect selection compared to signatures of direct selection. Additionally, evolutionary changes in recombination rates clearly impact signatures of indirect selection among species, which was not observable for signatures of direct selection. Finally, we provide evidence that signatures of indirect and direct selection do not show a strong association with each other and are driven by distinct evolutionary processes.

### Signatures of indirect selection are shaped by recombination rate dynamics

Speciation genomic studies across a wide range of species have revealed that differentiation islands tend to coincide with regions of low recombination as a consequence of a local reduction of genetic diversity due to indirect selection (21, 24–27, 30, 56). Despite the central role of recombination in shaping the differentiation landscape, the role of the evolutionary dynamics of recombination rate in these patterns has largely remained unexplored, often because recombination rate estimates have only been available for one of the studied species, or not available for the focal species, (21, 26, 27, 30, 56; but see 60).

Recent work has, however, demonstrated that not accounting for recombination rate evolution can lead to an underestimation of the impact of recombination rate on signatures of indirect selection (64), particularly in species where PRDM9 entails highly dynamic recombination rates such as mammals. Birds, in contrast, lack PRDM9 and their genomes are characterized by a more stable recombination landscape. Nevertheless, we are here able to demonstrate that evolutionary changes in recombination rate are central in driving differences in signatures of indirect selection between *Ficedula* flycatcher species, while evolutionarily stable reductions in recombination rate manifest in differentiation islands that are shared among species.

Characteristic for the *Ficedula* flycatchers is that both lineage-specific and shared signatures of indirect selection generally stretch several 100-kb, suggesting that the mechanism behind low-recombining regions acts at the broad-scale rather than the fine-scale. Chromosomal rearrangements are one such mechanism for broad-scale reduction in recombination rate (52, 53, 65). It is thus possible that a history of chromosomal rearrangements such as inversions between taiga flycatcher and collared flycatcher could explain lineage-specific differentiation islands and selective sweeps. In fact, the shared patterns of differentiation could potentially arise from a similar history. Although inversions are generally expected to reduce recombination when polymorphic within a species owing to the suppression of recombination in heterozygotes (66), there is evidence that reductions in recombination rate persist even after an inversion is fixed (67). In some cases, it also appears that shared inversions across populations have contributed to parallel evolution (68-70). While the role of inversions in the recombination dynamics of *Ficedula* flycatchers remains at present hypothetical, investigation of these hypotheses constitutes a relevant direction for future research that can benefit from recent advancements in long-read sequencing technology.

Yet, regardless of the molecular mechanism that drives local reductions in recombination rate, differentiation islands or selective sweeps would not arise without the action of indirect selection (12, 15-17). In addition, simulation studies suggest a relationship between the rate of selective sweeps and the adaptive substitution rate, at least in the absence of recombination (71). Since signatures of selective sweeps rapidly break down with increasing recombination rate (13, 14), the predicted relationship between the rate of selective sweeps and the adaptive substitution rate is less obvious in recombining genomes. Here, the *Ficedula* flycatcher system constitutes a relevant study system to explore this relationship, since the comparison of adaptive substitution rates in lineage-specific and shared *F*_*ST*_ peaks permits assessing the role of recombination and recombination dynamics. With variable recombination rate in lineage-specific *F*_*ST*_ peaks, we did not observe a relationship between lineage-specific *F*_*ST*_ peaks and any genomic signatures of direct selection. This may be because there has been no increase in adaptive substitutions in these regions or that any adaptive substitutions occurred too recently to be detected with the methods employed. Shared *F*_*ST*_ peaks and *F*_*ST*_ peaks coinciding with selective sweeps showed significantly higher π_N_/π_S_ for both species, which was driven by a greater reduction in π_S_ compared to π_N_ as both values were reduced compared to the background. Hitchhiking due to a selective sweep is known to cause a reduction in linked diversity (12), and theory suggests that the reduction in diversity is greater for synonymous sites than nonsynonymous sites (72). Although estimates of the adaptive substitution rate were elevated, in particular for taiga flycatcher, the increase was not significant for neither shared *F*_*ST*_ peaks nor *F*_*ST*_ peaks coinciding with selective sweeps compared to the genomic background. These results therefore indicate that differentiation islands and selective sweep signatures do not require significantly elevated rates of adaptation in order to manifest in the genome, and highlight again the central role of recombination dynamics in shaping indirect selection signatures.

### Recombination rate is not a major force in shaping signatures of direct selection

In addition to creating more pronounced signatures of indirect selection, tighter linkage between sites owing to low recombination rate is predicted to increase the impact of HRI. The consequence of this relationship is an increase in the efficacy of natural selection with increasing recombination rate, which has been observed in a number of systems (31, 34–37, 42). However, within both taiga flycatcher and collared flycatcher signatures of direct selection generally showed little variation with recombination rate, and did not reflect the relationships predicted by HRI.

There are several explanations why HRI may be less prevalent in *Ficedula* flycatchers compared to other systems. Selective interference occurs when multiple linked, selected mutations segregate within the population at the same time, and it has been demonstrated that the addition of neutral sites between selected sites may help to alleviate selective interference (43). This means that we are more likely to observe HRI where there is a greater density of functional sequences relative to the recombination rate. For example, gene density/cM in *D. melanogaster*, where evidence for HRI has been observed, is roughly an order of magnitude larger than in flycatcher (73; FlyBase release FB2022_04; Ensembl v. 107). Additionally, genetic diversity is much higher in *D. melanogaster* compared to flycatchers (74), meaning multiple selected mutations will be more likely to segregate at the same time. Taken together, these circumstances could explain why we find no effect of recombination rate on selection efficacy in the *Ficedula* flycatchers, even in the most gene dense regions of the genome. It is important to note, however, that evidence of HRI in birds is inconclusive (36, 38, 41, 75), and generalization among birds asks for further investigations.

To investigate if evolutionary changes in recombination rate could confound evidence of HRI in *Ficedula* flycatchers, we limited our analysis to genes where recombination rate appeared to be stable between taiga flycatcher and collared flycatcher. Still, no relationship between recombination rate and signatures of direct selection consistent with HRI appeared. Thus, recombination rate and its dynamics appear to not play a significant role in shaping any differences in signatures of direct selection between the two birds studied here. Lack of HRI in flycatchers is in fact in good agreement with earlier observations, which suggest that rather than the genomic background of a gene, functional characteristics of the gene such as its expression patterns are more likely to influence selection intensity on coding sequences (63). In line with this prediction, recent work found that estimates of *d*_N_/*d*_S_ were highly correlated among distantly related species with divergent genomic backgrounds (76), suggesting that conserved gene functions rather than genomic background shape the variation in *d*_N_/*d*_S_ among genes.

## Conclusions

By comparing recombination dynamics with genomic signatures of indirect and direct selection, we show that these different signatures of selection are not shaped by the same evolutionary processes. While signatures of indirect selection appear to be strongly shaped by the recombination landscape, signatures of direct selection are largely unaffected by the genomic background. Rather, gene function and expression patterns appear to play a more central role in shaping the efficacy of direct selection on protein-coding sequences.

## MATERIALS AND METHODS

### Samples and genotyping

Whole genome re-sequencing data for 187 flycatchers were collated from previously published work, including 65 taiga flycatchers (56), 15 red-breasted flycatchers (56), 95 collared flycatchers (57), 11 pied flycatchers (21), and one sample of snowy-browed flycatcher as an outgroup (21). BQSR recalibrated BAM files with reads mapped to the collared flycatcher reference genome FicAlb1.5 (60) were obtained for all samples. Genotyping was performed individually with HaplotypeCaller in GATK v.4.1, followed by GenotypeGVCFs with all samples combined specifying the *--all-sites* flag to genotype both variant and invariant sites. We removed indels and variable sites with more than one alternative allele using VCFtools v0.1.16 (77), resulting in a dataset containing 974,084,046 sites in total, including 116,929,990 SNVs.

Here, we focus only on sites located on the autosomes, excluding unassigned scaffolds, the Z-chromosome, and mitochondria. We applied hard filtering thresholds based on GATK best practices (QD < 2.0, FS > 60.0, MQ < 40.0, MQRankSum < -12.5, ReadPosRankSum < -8.0). We then applied genotype filters of GQ >30, DP >5 and DP <200. For the snowy-browed flycatcher, only the depth filters were applied (DP >5 and DP <200). Sites overlapping with annotated repeats for the collared flycatcher (78) were removed. We identified duplicated regions collapsed in the collared reference genome, using taiga flycatcher and collared flycatcher samples following previous work (56). After applying all genotype filters, we removed positions with more than 10% missing data in any of the four species. Our final dataset consisted of 566,724,393 callable sites including 51,424,863 SNVs.

### Ancestral sequence reconstruction and variant polarization

To polarize variable sites, we combined taiga and red-breasted flycatchers into one group and collared and pied flycatchers into another group, using Snowy-browed flycatcher as an outgroup. Whenever any two of the three groups were fixed for the same allele, this allele was considered the ancestral state. We reconstructed the ancestral reference genome by replacing respective sites of the collared flycatcher reference genome with the ancestral allele, and masking positions not genotyped in at least 90% of individuals in each species or where the ancestral allele was equivocal. This resulted in a set of 49,121,805 polarized SNVs.

### Estimation of recombination rate

Population-scaled recombination rate (*ρ* = 4N_e_r) was estimated for taiga flycatcher and collared flycatcher. Haplotypes were inferred with SHAPEIT2 (79), after removing singletons in both species. We adjusted the default settings of SHAPEIT2 to meet criteria for the collared flycatcher. Genome-wide rho was set to 0.037 (51) and effective population size was set to 300,000 (80). We set the --*states* parameter to 200 to improve the accuracy of phasing and *--window* to 0.5, which is recommended for WGS data. Phasing was performed separately for each scaffold, and two independent runs of phasing were performed for each scaffold in both species to validate robustness.

After phasing, recombination rate was estimated with LDhelmet (81) using the output from both runs of SHAPEIT2 separately. We provided prior probabilities of the ancestral allele frequency using the polarization data, which improves accuracy. The inferred ancestral state prior probability was set to 0.97, and the prior probability for each of the other three alleles to 0.01. For unpolarized sites, we set the prior probability to 0.25 for all four alleles. We provided a nucleotide substitution matrix from previous work in chicken (53). Because LDhelmet can handle a maximum of 50 haplotypes as input, we subset the phased data from both taiga flycatcher and collared flycatcher to the 25 unrelated samples with the least amount of missing data (Table S5). Following settings for *Ficedula* flycatchers established by (51), we ran five iterations of rjMCMC simulations for both species and both independent phased haplotypes to account for stochasticity in the recombination rate estimation. We set 2,000,000 iterations and 200,000 iterations of burn-in for each run, using a block penalty of 10 and a window of 50 SNPs.

We calculated the average recombination rate between SNP pairs for each of the 5 independent rjMCMC simulations and calculated the average LD recombination rate in 200-kb, 1-Mb, and 5-Mb genomic windows, weighted by the distance between SNP pairs accounting for SNP pairs spanning adjacent windows. To account for stochasticity in phasing, we performed a linear regression on the window-based estimates of rho from both phase outputs, and removed windows with standard residuals with an absolute value greater than 9 to exclude outliers. We then took the average value of rho across the two phases.

Window-based population-scaled recombination rates (*ρ* = 4*N*_*e*_*r*) were converted into cM/Mb by dividing by the genome-wide *N*_*e*_. Genome-wide *N*_*e*_ was estimated following (52), by fitting a robust linear regression without intercept between rho and the pedigree recombination map of collared flycatcher in 200-kb, 1-Mb, and 5-Mb windows (60) and obtaining the gradient of the fitted line between the two estimates of recombination rate. This gave an *N*_*e*_ estimate of 318,000 – 458,000 for taiga flycatcher, and 30,000 – 42,500 for the Gotland population of collared flycatcher, where the latter is in line with LD-based *N*_*e*_ estimates from (57). We used the value of *N*_*e*_ estimated from 5-Mb windows to convert estimates of recombination rate into cM/Mb, since this window size showed the strongest correlation with the linkage-map-based recombination rate (Table S6). After converting estimates, two outlier windows with much higher recombination rate (>20cM/Mb) in collared flycatcher were removed. In addition to window-based recombination rate estimates, we summarized the weighted average of recombination rate within each protein coding gene.

### Estimation of signatures of indirect selection within and between species

To assess signatures of indirect selection, we performed a selective sweep scan within taiga flycatcher and collared flycatcher separately. A composite likelihood ratio (CLR) test (82) was used to identify regions showing signatures of selective sweeps, using the polarized SNP data (see above) for the two species. The test was implemented in SweepFinder2 (83) using the -ug option with a pre-computed background SFS from each species (taiga flycatcher n = 65 diploid individuals; collared flycatcher n = 95), and a user-defined grid of locations for every variant. A significance threshold for the CLR test was based on simulations performed in SLiM 3, described previously (56), which incorporates background selection occurring across one 21 Mb chromosome using the collared flycatcher annotation (84) and recombination map (60) estimated from collared flycatcher chromosome 11. This gave a significance threshold of 46.25. We merged adjacent sites with significant CLR values to determine selective sweep regions, removing regions with only one significant site or with a site density less than 1/kb, and called presence/absence of selective sweeps in 200-kb genomic windows.

We also calculated window-based *F*_*ST*_ between the species pairs taiga and red-breasted flycatcher and collared and pied flycatcher, which reflects the history of linked selection between species. *F*_*ST*_ was calculated in 200-kb nonoverlapping windows using VCFtools v0.1.16 (77), which estimates Weir and Cockerham *F*_*ST*_ (85). To identify windows showing significantly elevated *F*_*ST*_ (so-called *F*_*ST*_ peaks), we Z-transformed *F*_*ST*_ separately for each chromosome and applied a Savitzky-Golay smoothing filter to the Z*F*_*ST*_ values using a polynomial of three and a filter length of seven. We defined *F*_*ST*_ peaks as windows with a smoothed Z*F*_*ST*_ value above 2 (i.e. two standard deviations above the chromosome mean). We categorized *F*_*ST*_ peaks as shared peaks if there was any overlap in the coordinates of an *F*_*ST*_ peak between the two species comparisons. *F*_*ST*_ peaks unique to taiga and red-breasted flycatcher were classified as taiga red-breasted unique (TR unique) and *F*_*ST*_ peaks unique to collared and pied flycatcher were classified as collared pied unique (CP unique). To compare recombination rate in different *F*_*ST*_ peak categories, we performed permutation test, randomizing the peak category and re-estimating the difference in mean recombination rate among 200-kb windows for 1,000 permutations.

### Multiple sequence alignments of protein-coding genes

To estimate signatures of direct selection on protein-coding sequences we constructed sequence alignments for one-to-one orthologues between taiga flycatcher, collared flycatcher, and zebra finch (*Taeniopygia guttata*). Zebra finch sequences were retrieved from Ensembl v. 104 for one-to-one orthologues between collared flycatcher and zebra finch. We reconstructed gene sequences for multiple sample sizes of taiga flycatcher and collared flycatcher, to account for the recent divergence time of the two species (approximately 3.76 coalescent units; 56) by masking polymorphic sites following the protocol outlined in (86).

For both species, we selected the 16 individuals (32 haplotypes) with the lowest amount of missing data (Table S5), and randomly down-sampled these individuals to obtain sample sizes of 1, 2, 8, 16, and 32 haplotypes. Gene sequences were reconstructed based on the collared flycatcher reference genome, after masking non-callable sites from the variant calling dataset described above. Within each sample size, we masked polymorphic sites and replaced the reference allele with the alternative allele for sites where the alternative allele was fixed in the respective species. We removed genes from further analysis if more than 50% of sites were masked in taiga flycatcher or collared flycatcher, which resulted in a set of 7052 genes.

For each gene we combined the sequences for all samples into a single fasta file and aligned the sequences using prank v170427 (87). Prior to alignment with prank, we ran clustalw v2.1 (88) to estimate a guide tree for each gene, which we then provided as input to prank, running prank with the *-once* flag to perform one iteration.

### Estimation of signatures of direct selection within and between species

We estimated signatures of direct selection in taiga flycatcher and collared flycatcher for one-to-one orthologues between flycatcher and zebra finch that passed filtering thresholds described above. The polymorphism-based measure of direct selection, π_N_/π_S_, was calculated based on four-fold and zero-fold degenerate sites for taiga flycatcher (n=65) and collared flycatcher (n=95). Four-fold and zero-fold degenerate sites were extracted from the reconstructed ancestral genome with a custom python script (https://github.com/madeline-chase/flycatcher_recom). We subset the variable four-fold and zero-fold degenerate sites for only GC-conservative variants, i.e. strong-to-strong (C&G) variants or weak-to-weak (A&T) variants, to correct estimates of π_N_/π_S_ for any potential influence of gBGC. π_S_ and π_N_ were then calculated from allele frequency data following (56), where the per site estimate of π is obtained by dividing by the total number of four-fold or zero-fold sites.

As another measure of direct selection, we estimated *d*_N_/*d*_S_ for taiga flycatcher and collared flycatcher using Bio++ v3.0 (89). We estimated the gene tree for each gene separately using a strand-symmetric L95 model (90) implemented in bppml, and then mapped substitutions to the tree using mapnh, as implemented in (91). This approach allowed us to separate substitutions into different categories and correct for gBGC by restricting the analysis to only GC-conservative changes. To retrieve an estimate of *d*_N_/*d*_S_ for each gene, we weighted the counts of nonsynonymous and synonymous substitutions by the proportion of GCs or ATs for strong-to-strong and weak-to-weak variants, respectively. We excluded genes with an exon length below 200 and with norm values above 10 for the GC-conservative estimates. We summarized *d*_N_/*d*_S_, by dividing the average value of *d*_N_ by the average value of *d*_S_. There was little evidence of sample size dependence in the estimates of *d*_N_/*d*_S_ (Table S7; Table S8), suggesting that divergence between taiga flycatcher and collared flycatcher is deep enough for phylogenetic analysis (86). All further analyses with *d*_N_/*d*_S_ were therefore only based on the 32 haplotype sample size for both taiga flycatcher and collared flycatcher.

Using the four-fold and zero-fold site frequency spectra and the results of *d*_N_/*d*_S_ for both taiga flycatcher and collared flycatcher, we estimated the adaptive substitution rate (ω_a_) with the software DFE-alpha (92, 93). For both species, we compared the 1-epoch and 2-epoch model of demographic history based on all genes using a likelihood ratio test, and found the 2-epoch model was a better fit (Table S9; Table S10), which we then used for all further analysis.

### Gene binning and permutation tests for differences among bins

We divided the gene-based estimates of recombination rate for both taiga flycatcher and collared flycatcher into three quantiles of approximately equal numbers of genes to obtain protein-coding genes with low (L), intermediate (M), and high (H) recombination rate in order to compare measures of direct selection in regions of varying recombination rate. To investigate the effect of changes in the recombination landscape on measures of direct selection, we subset the three recombination rate bins for genes showing conserved recombination rate between taiga flycatcher and collared flycatcher. For this purpose, we performed a principal component analysis (PCA) using gene-based recombination rate estimates for taiga flycatcher and collared flycatcher and identified genes with conserved recombination rate if they were within one standard deviation from the mean of the second principal component. Finally, we estimated functional sequence density in 200-kb windows, and divided windows into three approximately equal sized quantiles of low, intermediate and high functional density and identified genes overlapping with either low or high functional density windows for each of the three recombination rate bins. We then estimated π_N_/π_S_, *N*_e_*s, d*_N_/*d*_S_, and ω_a_ for genes in each set of recombination rate bins (all genes, conserved recombination rate, low gene density, and high gene density), for all changes and for GC-conservative changes, respectively.

We tested for a statistically significant difference between measures of selection estimated in different recombination rate bins using a permutation test. For this purpose, we re-shuffled the genes among the three recombination rate bins, keeping the bin sizes the same. We calculated the randomized difference between each comparison of recombination rate, and computed the *p*-value based on the number of permutations that showed a difference as large or larger than the observed difference between those bins, out of 1,000 permutations.

We used a similar permutation test approach to test for statistically significant differences in signatures of selection for *F*_*ST*_ peaks, where we compared each *F*_*ST*_ peak category to the background of genes not overlapping *F*_*ST*_ peaks, randomized the genes found in both, and compared our permuted differences to the observed values. We again performed 1,000 permutations.

## Data accessibility

Sequencing data for all samples are available at the EMBL-EBI European Nucleotide Archive (ENA; http://www.ebi.ac.uk/ena) with the following accession numbers: PRJEB43825 (taiga and red-breasted flycatchers), PRJEB22864 (collared flycatcher), and PRJEB7359 (pied and snowy-browed flycatchers). VCF files used for analyses will be deposited on dryad after publication. Scripts used for analysis are available at https://github.com/madeline-chase/flycatcher_recom.

## Acknowledgements

We thank Laurent Guéguen for generous advice on running Bio++, and Laurent Duret for helpful feedback on an earlier version of this manuscript. This study was funded by grants from the Swedish Research Council (2013-8271 to Hans Ellegren), and the Knut and Alice Wallenberg Foundation (2014/0044 to Hans Ellegren). Computations were performed on resources provided by the Swedish National Infrastructure for Computing (SNIC) through Uppsala Multidisciplinary Center for Advanced Computational Science (UPPMAX).

## Author contributions

MAC and CFM conceived of the study. CFM supervised the study. MAC performed the data analyses. MAC and CFM wrote the manuscript and approved the final version.

## Notes

### Competing Interest Statement

The authors have declared no competing interest.

