## Supplemental Material for "The role of recombination dynamics in shaping signatures of direct and indirect selection across the *Ficedula* flycatcher genome"

### Supplemental tables

**Table S1:** Number of 200-kb windows in each $F_{ST}$ peak category

**Table S2:** Number of genes in recombination rate bins, for all genes and genes with conserved recombination rate

**Table S3:** Number of genes in recombination rate bins, for genes in low functional density regions and genes in high functional density regions

**Table S4:** Estimates of direct selection within $F_{ST}$ peaks for all changes

**Table S5:** List of samples genotyped with proportion of missing data

**Table S6:** Correlation analysis between window-based rho and pedigree map for taiga flycatcher and collared flycatcher

**Table S7:** Low sample-size dependence in $d_N/d_S$ for GC-conservative changes

**Table S8:** Low sample-size dependence in $d_N/d_S$ for all changes

**Table S9:** DFE-alpha estimates for 1-epoch vs. 2-epoch models for GC-conservative changes

**Table S10:** DFE-alpha estimates for 1-epoch vs. 2-epoch models for all changes

### Supplemental figures

**Figure S1:** The correlation of recombination rate between species

**Figure S2:** The relationship between selective sweeps and recombination rate

**Figure S3:** The relationship between $F_{ST}$ and recombination rate

**Figure S4:** The impact of recombination on signatures of direct selection for all substitutions

**Figure S5:** The impact of gene density on signatures of direct selection for all substitutions.

**Table S1:** Number of 200-kb windows in each of four $F_{ST}$ peak categories, including No peak, Shared peak, TR (taiga/red-breasted) unique, and CP (collared/pied) unique.

| Peak category | Number of windows |
|---|---|
| No peak | 4224 |
| Shared peak | 132 |
| TR unique | 114 |
| CP unique | 80 |

**Table S2:** Number of genes in recombination rate bins for taiga flycatcher and collared flycatcher, split by all genes and genes with conserved recombination rate. The number of genes are given for GC-conservative analyses, with genes used for the all-sites analysis given in parentheses. Differences in gene number result from filtering of outlier results.

| Recombination bin | Taiga flycatcher | Collared flycatcher |
|---|---|---|
| Low (L), all | 2228 (2244) | 2152 (2174) |
| Low (L), conserved | 1787 (1803) | 1781 (1797) |
| Intermediate (M), all | 2280 (2301) | 2308 (2339) |
| Intermediate (M), conserved | 2039 (2059) | 2019 (2046) |
| High (H), all | 1921 (1931) | 1951 (1962) |
| High (H), conserved | 1230 (1236) | 1246 (1254) |

**Table S3:** Number of genes in recombination rate bins for taiga flycatcher and collared flycatcher, split by genes in low functional density regions and high functional density regions. The number of genes are given for GC-conservative analyses, with genes used for the all-sites analysis given in parentheses. Differences in gene number result from filtering of outlier results.

| Recombination bin | Taiga flycatcher | Collared flycatcher |
|---|---|---|
| Low (L), low density | 263 (264) | 212 (214) |
| Low (L), high density | 901 (905) | 1012 (1021) |
| Intermediate (M), low density | 239 (244) | 235 (236) |
| Intermediate (M), high density | 970 (979) | 1029 (1039) |
| High (H), low density | 107 (108) | 165 (166) |
| High (H), high density | 1142 (1148) | 967 (972) |

**Table S4**: Comparison of indirect and direct selection, all changes. Shown are estimates of the efficacy of direct selection for genes grouped by $F_{ST}$ peak category. Categories include genes outside $F_{ST}$ peaks, genes in $F_{ST}$ peaks shared between species comparisons, genes within peaks specific to the focal species, genes within peaks that show a signature of a selective sweep, and genes within peaks that do not show a signature of a selective sweep. *p*-values comparing whether statistics within $F_{ST}$ peaks are significantly different from outside $F_{ST}$ peaks, are presented in parentheses. Significant differences are indicated in bold, *p*-value < 0.05

| | **Taiga flycatcher** | | | | | **Collared flycatcher** | | | | |
|---|---|---|---|---|---|---|---|---|---|---|
| | $\pi_N/\pi_S$ | $N_e s$ | $d_N/d_S$ | $\omega_a$ | **genes** | $\pi_N/\pi_S$ | $N_e s$ | $d_N/d_S$ | $\omega_a$ | **genes** |
| **Outside peaks** | 0.18 | 845 | 0.17 | 0.073 | 5976 | 0.19 | 3711 | 0.16 | 0.0020 | 5975 |
| **Shared peaks** | **0.25 (0.005)** | **167 (0.020)** | **0.23 (0.013)** | **0.15 (0.001)** | 223 | **0.26 (0.004)** | 421 (0.25) | **0.22 (0.006)** | 0.045 (0.067) | 223 |
| **Lineage specific peaks** | 0.20 (0.24) | **227 (0.027)** | 0.17 (0.99) | **0.12 (0.046)** | 197 | 0.20 (0.80) | 257 (0.30) | 0.15 (0.87) | 0.047 (0.18) | 98 |
| **Peaks with sweep overlap** | **0.23 (0.002)** | **155 (0.005)** | **0.21 (0.048)** | **0.15 (0.001)** | 324 | **0.27 (0.002)** | 334 (0.25) | 0.19 (0.12) | 0.016 (0.55) | 213 |
| **Peaks with no sweep overlap** | 0.24 (0.068) | 322 (0.14) | 0.19 (0.43) | 0.11 (0.23) | 96 | 0.20 (0.85) | 399 (0.34) | **0.22 (0.038)** | **0.10 (0.004)** | 108 |

**Table S5:** Samples genotyped for taiga flycatcher and collared flycatcher, and proportion of variable sites with missing data for each sample. Samples included in LDhelmet analysis are marked with * and samples included for polymorphism-aware $d_N/d_S$ estimation are marked with †.

| Sample | Species | Missing data |
|---|---|---|
| Sample_1 | Taiga | 0.03660 |
| Sample_10*† | Taiga | 0.00050 |
| Sample_11* | Taiga | 0.00080 |
| Sample_12* | Taiga | 0.00058 |
| Sample_13 | Taiga | 0.11069 |
| Sample_16 | Taiga | 0.00368 |
| Sample_17 | Taiga | 0.00551 |
| Sample_18*† | Taiga | 0.00055 |
| Sample_2 | Taiga | 0.00140 |
| Sample_20*† | Taiga | 0.00057 |
| Sample_21* | Taiga | 0.00059 |
| Sample_22 | Taiga | 0.00750 |
| Sample_23 | Taiga | 0.00214 |
| Sample_24* | Taiga | 0.00118 |
| Sample_25 | Taiga | 0.00310 |
| Sample_26 | Taiga | 0.00108 |
| Sample_28 | Taiga | 0.00490 |
| Sample_29 | Taiga | 0.00399 |
| Sample_3 | Taiga | 0.01705 |
| Sample_30 | Taiga | 0.00124 |
| Sample_33 | Taiga | 0.00091 |
| Sample_34 | Taiga | 0.00291 |
| Sample_35 | Taiga | 0.02405 |
| Sample_36 | Taiga | 0.00160 |
| Sample_37* | Taiga | 0.00079 |
| Sample_38*† | Taiga | 0.00048 |
| Sample_39*† | Taiga | 0.00059 |
| Sample_40 | Taiga | 0.00393 |
| Sample_41* | Taiga | 0.00123 |
| Sample_42* | Taiga | 0.00122 |
| Sample_44 | Taiga | 0.01342 |
| Sample_45 | Taiga | 0.00144 |
| Sample_5 | Taiga | 0.01830 |
| Sample_57*† | Taiga | 0.00070 |
| Sample_58 | Taiga | 0.00402 |

| | | |
|---|---|---|
| Sample_59*† | Taiga | 0.00046 |
| Sample_6 | Taiga | 0.01074 |
| Sample_60*† | Taiga | 0.00034 |
| Sample_61*† | Taiga | 0.00033 |
| Sample_63 | Taiga | 0.09270 |
| Sample_64*† | Taiga | 0.00036 |
| Sample_66* | Taiga | 0.00122 |
| Sample_67 | Taiga | 0.01742 |
| Sample_68 | Taiga | 0.00150 |
| Sample_69 | Taiga | 0.07011 |
| Sample_7 | Taiga | 0.00237 |
| Sample_70 | Taiga | 0.00154 |
| Sample_71*† | Taiga | 0.00035 |
| Sample_72 | Taiga | 0.00135 |
| Sample_73*† | Taiga | 0.00034 |
| Sample_74 | Taiga | 0.00104 |
| Sample_75 | Taiga | 0.00344 |
| Sample_76 | Taiga | 0.00761 |
| Sample_78 | Taiga | 0.01162 |
| Sample_79*† | Taiga | 0.00043 |
| Sample_8 | Taiga | 0.07827 |
| Sample_80*† | Taiga | 0.00056 |
| Sample_81*† | Taiga | 0.00034 |
| Sample_86 | Taiga | 0.00176 |
| Sample_87 | Taiga | 0.00185 |
| Sample_88* | Taiga | 0.00083 |
| Sample_89 | Taiga | 0.02787 |
| Sample_9 | Taiga | 0.00080 |
| Sample_90 | Taiga | 0.00171 |
| Sample_91*† | Taiga | 0.00031 |
| Sample_15F129 | Collared | 0.00041 |
| Sample_15F130 | Collared | 0.00121 |
| Sample_15F131 | Collared | 0.00038 |
| Sample_15F135 | Collared | 0.00072 |
| Sample_15F142 | Collared | 0.00083 |
| Sample_15F143 | Collared | 0.00115 |
| Sample_15F145 | Collared | 0.00045 |
| Sample_15F149 | Collared | 0.00054 |
| Sample_15F151 | Collared | 0.00048 |
| Sample_15F17 | Collared | 0.00059 |
| Sample_15F18* | Collared | 0.00029 |
| Sample_15F21 | Collared | 0.00075 |

| | | |
|---|---|---|
| Sample_15F22 | Collared | 0.00052 |
| Sample_15F23 | Collared | 0.00075 |
| Sample_15F24 | Collared | 0.00049 |
| Sample_15F25 | Collared | 0.00077 |
| Sample_15F29* | Collared | 0.00033 |
| Sample_15F447 | Collared | 0.00070 |
| Sample_15F448*† | Collared | 0.00021 |
| Sample_15F450*† | Collared | 0.00020 |
| Sample_15F453* | Collared | 0.00206 |
| Sample_15F457 | Collared | 0.00160 |
| Sample_15F459 | Collared | 0.00055 |
| Sample_15F460 | Collared | 0.00042 |
| Sample_15M153 | Collared | 0.00048 |
| Sample_15M155 | Collared | 0.00547 |
| Sample_15M158 | Collared | 0.01165 |
| Sample_15M160 | Collared | 0.00117 |
| Sample_15M161* | Collared | 0.00052 |
| Sample_15M162 | Collared | 0.00070 |
| Sample_15M163 | Collared | 0.00200 |
| Sample_15M201 | Collared | 0.00195 |
| Sample_15M202 | Collared | 0.00337 |
| Sample_15M203 | Collared | 0.00072 |
| Sample_15M204* | Collared | 0.00046 |
| Sample_15M207 | Collared | 0.00187 |
| Sample_15M468 | Collared | 0.00078 |
| Sample_15M469 | Collared | 0.00032 |
| Sample_15M475 | Collared | 0.00215 |
| Sample_15M477 | Collared | 0.00052 |
| Sample_15M49 | Collared | 0.00120 |
| Sample_15M537 | Collared | 0.00080 |
| Sample_15M568* | Collared | 0.00046 |
| Sample_15M571 | Collared | 0.00081 |
| Sample_15M573*† | Collared | 0.00031 |
| Sample_15M589 | Collared | 0.00213 |
| Sample_15M684*† | Collared | 0.00019 |
| Sample_15M724 | Collared | 0.00289 |
| Sample_93F24 | Collared | 0.00068 |
| Sample_93F26 | Collared | 0.00046 |
| Sample_93F30* | Collared | 0.00033 |
| Sample_93F32 | Collared | 0.00082 |
| Sample_93F34 | Collared | 0.00039 |
| Sample_93F35 | Collared | 0.00085 |

| | | |
|---|---|---|
| Sample_93F42*† | Collared | 0.00028 |
| Sample_93F44 | Collared | 0.00044 |
| Sample_93F45*† | Collared | 0.00016 |
| Sample_93F47 | Collared | 0.00198 |
| Sample_93F54*† | Collared | 0.00026 |
| Sample_93F56 | Collared | 0.00086 |
| Sample_93F59*† | Collared | 0.00027 |
| Sample_93F74*† | Collared | 0.00029 |
| Sample_93F75*† | Collared | 0.00024 |
| Sample_93F77 | Collared | 0.00085 |
| Sample_93F82 | Collared | 0.00094 |
| Sample_93F88 | Collared | 0.00055 |
| Sample_93F89*† | Collared | 0.00025 |
| Sample_93F90 | Collared | 0.00037 |
| Sample_93F92 | Collared | 0.00038 |
| Sample_93F93 | Collared | 0.00072 |
| Sample_93F94 | Collared | 0.00041 |
| Sample_93M25*† | Collared | 0.00022 |
| Sample_93M27 | Collared | 0.00193 |
| Sample_93M28 | Collared | 0.00081 |
| Sample_93M29 | Collared | 0.00059 |
| Sample_93M36 | Collared | 0.00104 |
| Sample_93M38*† | Collared | 0.00042 |
| Sample_93M39 | Collared | 0.00070 |
| Sample_93M40*† | Collared | 0.00036 |
| Sample_93M41 | Collared | 0.00082 |
| Sample_93M46*† | Collared | 0.00024 |
| Sample_93M53 | Collared | 0.00206 |
| Sample_93M55 | Collared | 0.00057 |
| Sample_93M58 | Collared | 0.00046 |
| Sample_93M71 | Collared | 0.00066 |
| Sample_93M72 | Collared | 0.00090 |
| Sample_93M73 | Collared | 0.00066 |
| Sample_93M78 | Collared | 0.00060 |
| Sample_93M79* | Collared | 0.00046 |
| Sample_93M80 | Collared | 0.00150 |
| Sample_93M81*† | Collared | 0.00019 |
| Sample_93M83 | Collared | 0.00207 |
| Sample_93M84 | Collared | 0.00076 |
| Sample_93M86* | Collared | 0.00046 |
| Sample_93M91 | Collared | 0.00053 |

**Table S6:** Correlation analysis between window-based rho for taiga flycatcher and collared flycatcher and pedigree-based recombination rate for collared flycatcher for three genomic window sizes.

| Species | 200-kb windows | 1-Mb windows | 5-Mb windows |
|---|---|---|---|
| **Taiga** | 0.46 | 0.55 | 0.77 |
| **Collared** | 0.48 | 0.53 | 0.76 |

**Table S7:** Low sample-size dependence in $d_N/d_S$ for GC-conservative changes.

| Sample size | $d_N$ | $d_S$ | $d_N/d_S$ |
|---|---|---|---|
| **Taiga 1** | 0.20 | 1.0 | 0.20 |
| **Taiga 2** | 0.20 | 0.99 | 0.20 |
| **Taiga 8** | 0.20 | 1.0 | 0.20 |
| **Taiga 16** | 0.20 | 0.99 | 0.20 |
| **Taiga 32** | 0.21 | 1.0 | 0.21 |
| **Collared 1** | 0.18 | 0.95 | 0.19 |
| **Collared 2** | 0.17 | 0.93 | 0.19 |
| **Collared 8** | 0.18 | 0.87 | 0.20 |
| **Collared 16** | 0.18 | 0.91 | 0.20 |
| **Collared 32** | 0.18 | 0.90 | 0.20 |

**Table S8:** Low sample-size dependence in $d_N/d_S$ for all changes.

| Sample size | dN | dS | $d_N/d_S$ |
|---|---|---|---|
| **Taiga 1** | 0.41 | 2.5 | 0.16 |
| **Taiga 2** | 0.42 | 2.5 | 0.16 |
| **Taiga 8** | 0.42 | 2.5 | 0.17 |
| **Taiga 16** | 0.42 | 2.5 | 0.17 |
| **Taiga 32** | 0.43 | 2.5 | 0.17 |
| **Collared 1** | 0.40 | 2.6 | 0.15 |
| **Collared 2** | 0.39 | 2.6 | 0.15 |
| **Collared 8** | 0.40 | 2.6 | 0.16 |
| **Collared 16** | 0.40 | 2.6 | 0.16 |
| **Collared 32** | 0.41 | 2.5 | 0.16 |

**Table S9:** DFE-alpha estimates for 1-epoch vs. 2-epoch models GC-conservative changes.

| | **1-epoch** | | **2-epoch** | | | | | |
|---|---|---|---|---|---|---|---|---|
| **Species** | **N1** | **L** | **N1** | **N2** | **T2** | **L** | **LRT** | ***p*-value** |
| **Taiga** | 100 | -83003 | 100 | 879 | 121 | -82655 | 696 | $< 2.2 \cdot 10^{-16}$ |
| **Collared** | 100 | -37115 | 100 | 50 | 500 | -37111 | 8.28 | $0.16 \cdot 10^{-2}$ |

**Table S10:** DFE-alpha estimates for 1-epoch vs. 2-epoch models for all changes.

| | 1-epoch | | 2-epoch | | | | | |
|---|---|---|---|---|---|---|---|---|
| **Species** | **N1** | **L** | **N1** | **N2** | **T2** | **L** | **LRT** | ***p*-value** |
| **Taiga** | 100 | -442539 | 100 | 1000 | 175 | -440007 | 5065 | $< 2.2 \cdot 10^{-16}$ |
| **Collared** | 100 | -210473 | 100 | 55 | 550 | -210383 | 180 | $< 2.2 \cdot 10^{-16}$ |

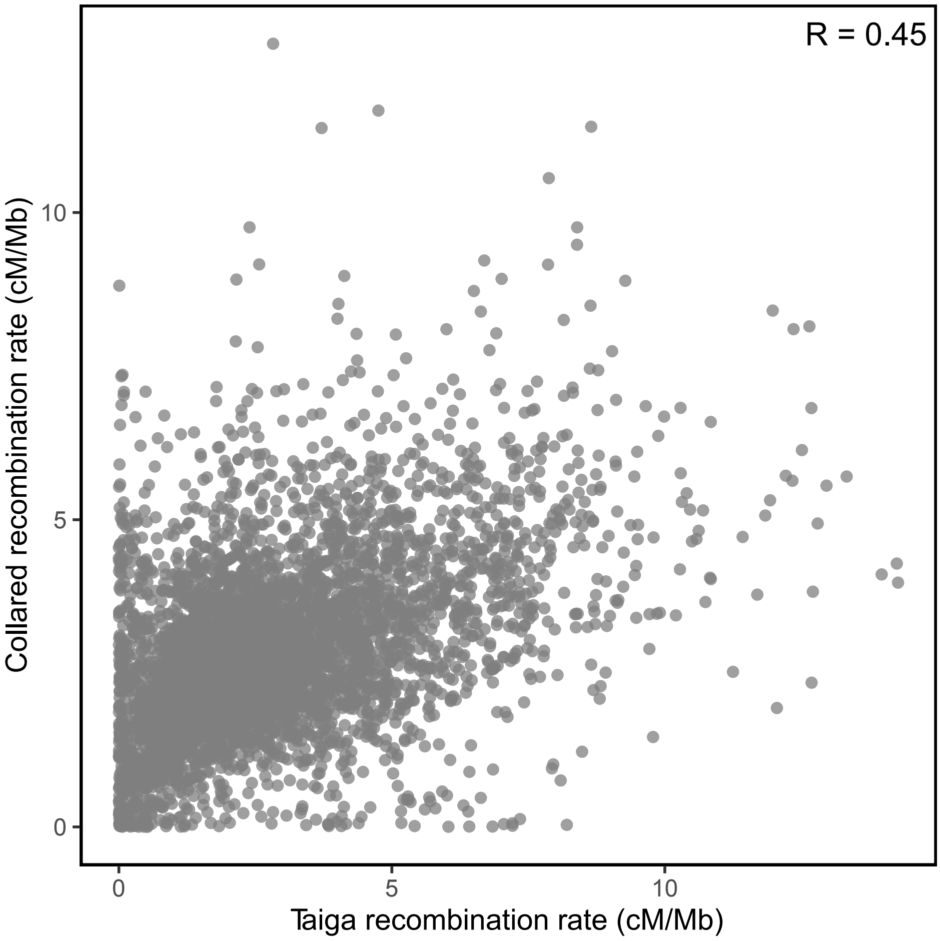


**Figure S1**: Correlation of LD-based recombination rate between taiga flycatcher and collared flycatcher, estimated in 200-kb genomic windows.

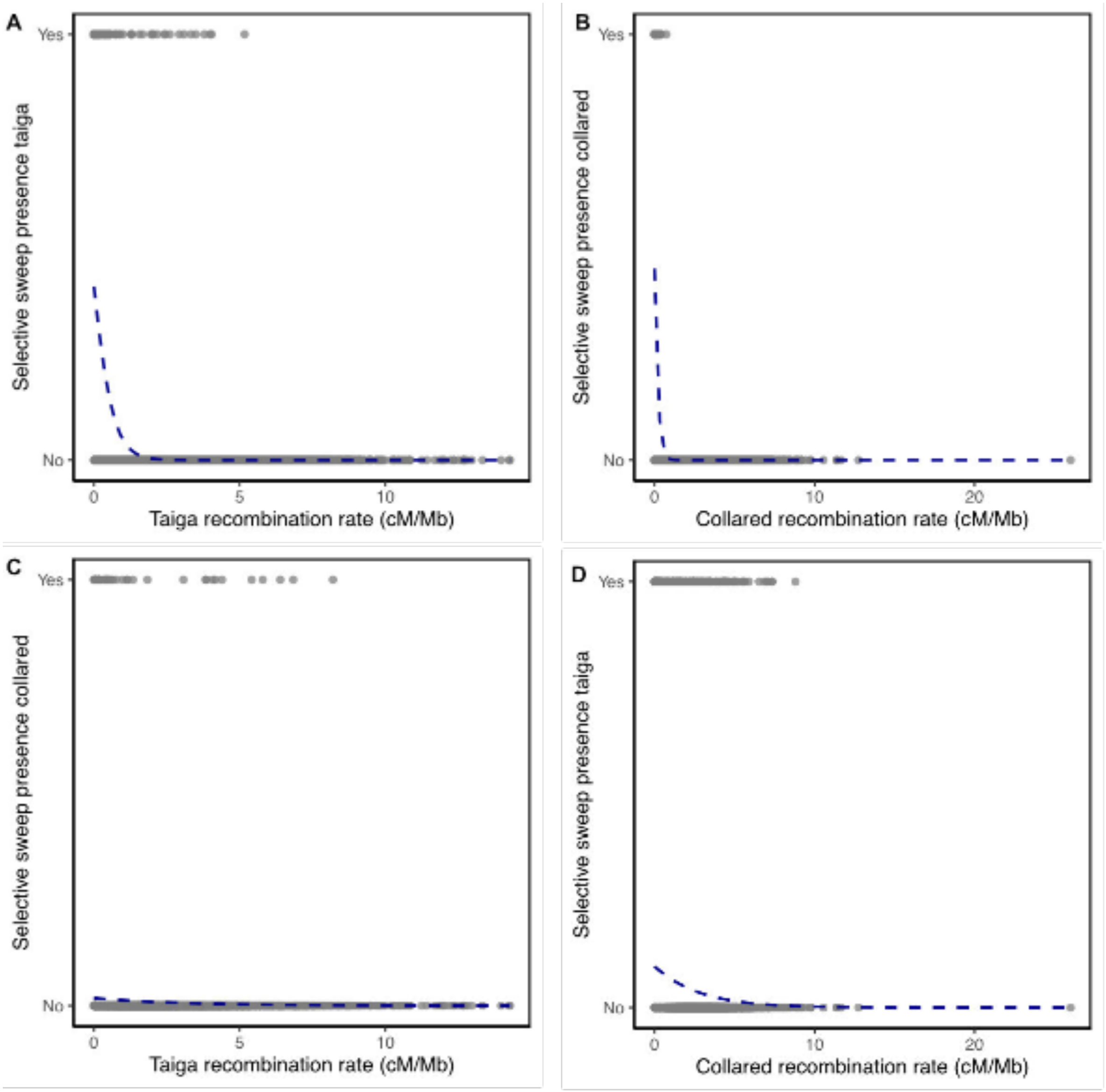


**Figure S2:** Relationship between selective sweeps and recombination rate. Shown is presence or absence of selective sweeps in 200-kb genomic windows for both taiga flycatcher and collared flycatcher, compared to recombination rate estimates for both species. The dashed blue line shows the fitted logistic regression.

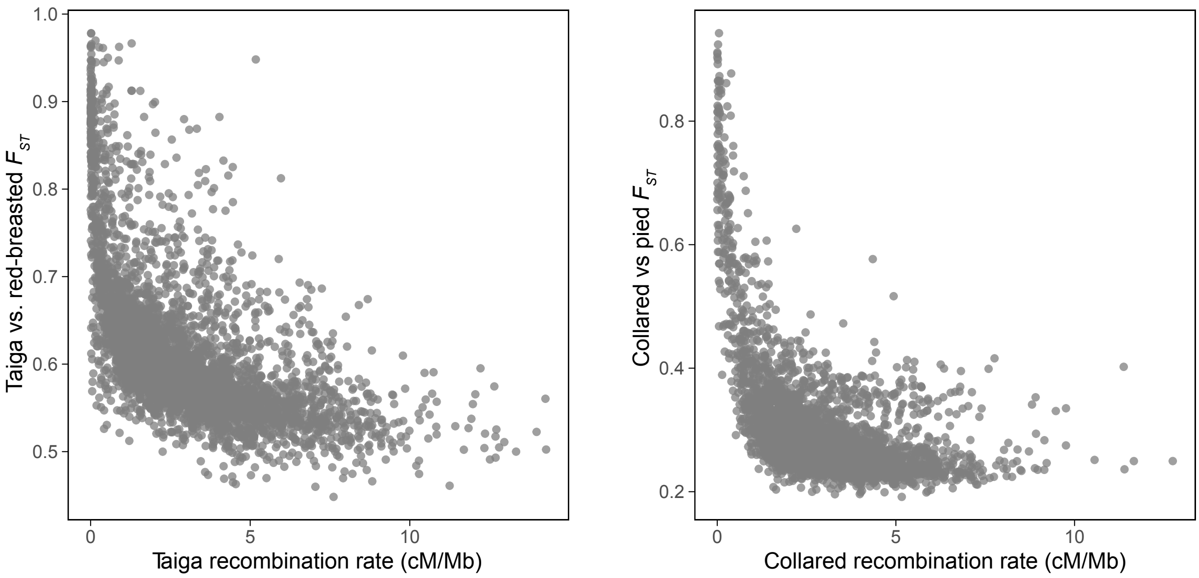


**Figure S3:** Relationship between $F_{ST}$ and recombination rate estimated in 200-kb genomic windows.

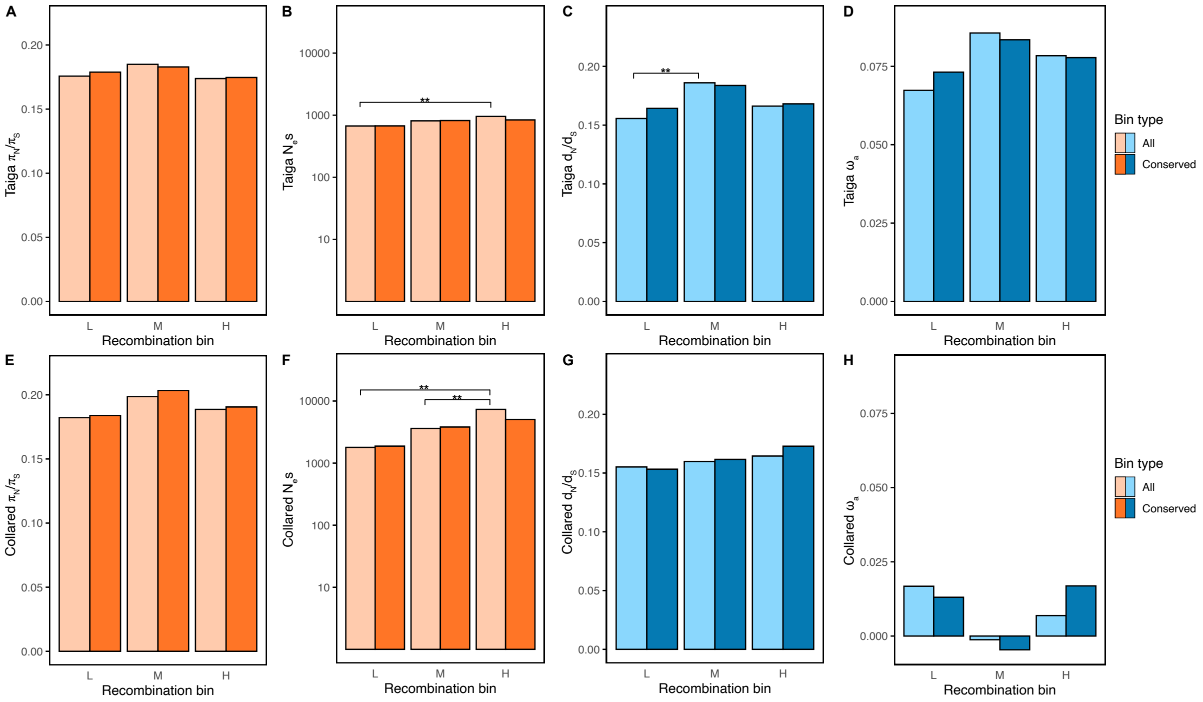


**Figure S4: Impact of recombination rate on signatures of direct selection for all substitutions.** Shown are estimates of $\pi_N/\pi_S$, $N_e s$, $d_N/d_S$ and $\omega_a$ for (A-D) taiga flycatcher and for (E-H) collared flycatcher estimated for genes overlapping with 3 bins of recombination rate (low (L), intermediate (M) and high (H)). Genes are separated based on conservation of recombination rate in taiga flycatcher and collared flycatcher. Colours for bin type correspond to whether the estimate shown is calculated for all genes (lighter) or for genes overlapping with windows with conserved recombination rate (darker). Tests for significance were performed between recombination rate within bin types (all or conserved). Asterisks denote statistical significance, * p <0.05; ** p<0.01. See Table S2 for the number of genes in each bin.

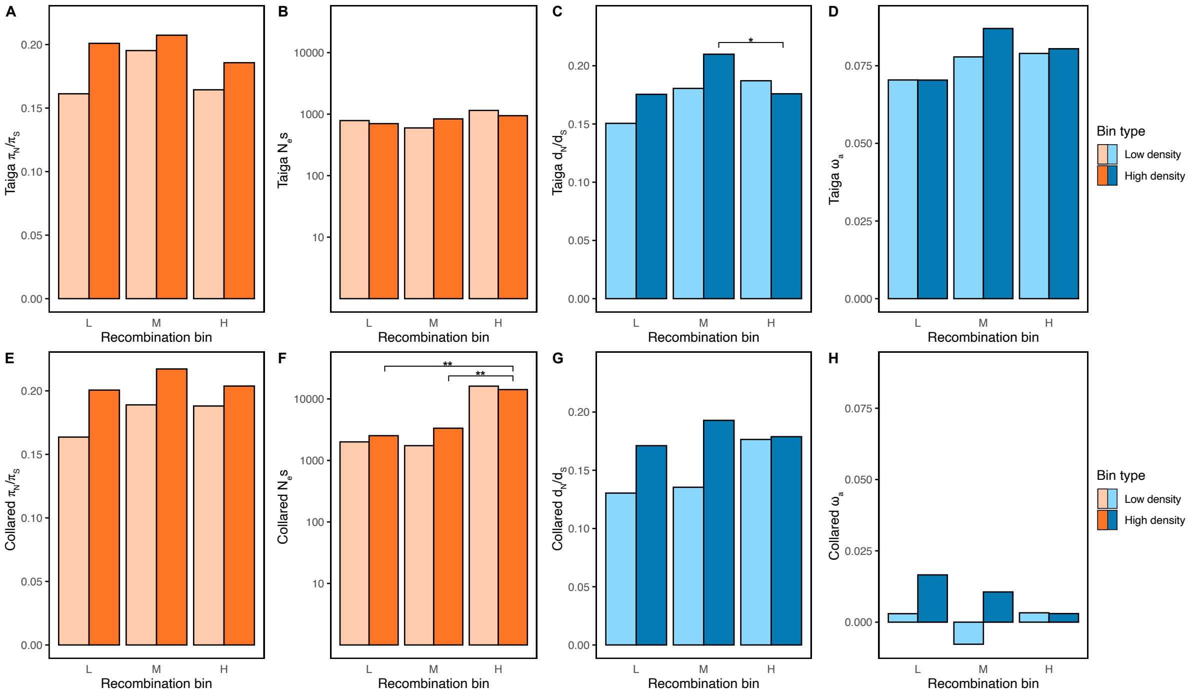


**Figure S5: The impact of gene density on signatures of direct selection for all substitutions.** Shown are estimates of $\pi_N/\pi_S$, $N_e s$, $d_N/d_S$ and $\omega_a$ for (A-D) taiga flycatcher and for (E-H) collared flycatcher estimated for genes overlapping with 3 bins of recombination rate (low (L), intermediate (M) and high (H)). Genes are separated based on whether they are found in windows with low or high density of functional sites. Tests for significance were performed between recombination bins within both bin density types. Asterisks denote statistical significance based on 1000 permutations, * $p < 0.05$, ** $p < 0.01$. See Table S3 for the number of genes in each bin.
